# The B chromosome of *Pseudococcus viburni:* a selfish chromosome that exploits whole-genome meiotic drive

**DOI:** 10.1101/2021.08.30.458195

**Authors:** Isabelle M. Vea, Andrés G. de la Filia, Kamil S. Jaron, Andrew J. Mongue, Francisco J. Ruiz-Ruano, Scott E.J Barlow, Ross Nelson, Laura Ross

## Abstract

Meiosis, the key process underlying sexual reproduction, is generally a fair process: each chromosome has a 50% chance of being included into each gamete. However in some organisms meiosis has become highly aberrant with some chromosomes having a higher chance of making it into gametes than others. Yet why and how such systems evolve remains unclear. Here we study the unusual reproductive genetics of mealybugs, in which only maternal-origin chromosomes are included into the gametes during male meiosis, while paternally-derived chromosomes degrade. This “whole genome meiotic drive” occurs in all males and is evolutionarily conserved. However one species - the obscure mealybug *Pseudococcus viburni* - has a segregating B chromosome that increases in frequency by escaping paternal genome elimination. Here we present whole-genome and gene expression data from laboratory lines with and without B chromosomes. These data allow us to identify B-linked sequences including >70 protein-coding genes as well as a B-specific satellite repeat that makes up a significant proportion of the chromosome. We also used these data to investigate the evolutionary origin of the B chromosome. The few paralogs between the B and the core genome are distributed throughout the genome, showing that it is unlikely that the B originated through a simple duplication of one of the autosomes. We also find that while many of the B-linked genes do not have paralogs within the *P*.*viburni* genome, but they do show orthology with genes in other hemipteran insects suggesting that the B might have originated from fission of one of the autosomes, possibly followed by further translocations of individual genes. Finally in order to understand the mechanisms by which the B is able to escape elimination when paternally-derived we generated gene expression data for males and females with and without B chromosomes. We find that at the developmental stage when meiosis is taking place only a small number of B-linked genes show significant expression. Only one gene was significantly over-expressed during male meiosis, which is when the drive occurs: a acetyltransferase involved in H3K56Ac, which has a putative role in meiosis and is therefore a promising candidate for further studies. Together, these results form a promising foundation for studying the mechanisms of meiotic drive in a system that is uniquely suited for this approach.

## Introduction

Multicellular organisms can be regarded as nested hierarchies of cooperating entities: genes within chromosomes, chromosomes within cells, and cells within individuals. But under the surface there is a hidden world of evolutionary conflict. The integrity of higher levels is challenged by lower-level entities trying to improve their representation in future generations (Hurst 1992; Szathmáry and Smith 1995; Burt and Trivers 2006). Intragenomic conflict – which arises when different entities within a genome have antagonic evolutionary agendas (Gardner and Úbeda 2017) – is especially pronounced during reproduction, since only limited numbers of genes are transmitted to the next generation. Meiotic drive – whereby a selfish genetic element hijacks gametogenesis to increase its transmission to future generations – is a key mechanism for intragenomic conflict (Lyttle 1993; Lindholm et al. 2016; Lenormand et al. 2016). Normally, during meiosis, each gene has a 50% chance of being passed on to the gametes. Meiotic drivers however can bias this fair gamble, increasing their transmission to the offspring at the expense of other genes. Which part of the genome “drives” can vary from just a single gene, to individual chromosomes, and even in some cases entire haploid genomes (reviewed in Burt and Trivers 2006; Gardner and Ross 2014). However, despite an abundance of drive systems across the tree of life, we know very little about their origin, evolution, or the mechanisms that these drivers use to increase their transmission.

One of the most widespread classes of meiotic drivers are supernumerary B chromosomes. They are found across fungi, plants and animals and estimated to be present in at least 10% of eukaryotic species (Camacho 2005; Ahmad and Martins 2019). B chromosomes are genomic parasites that exist within cells of some individuals of a species, sometimes in multiple copies, in addition to autosomes and sex chromosomes – the A chromosomes (Camacho et al. 2000; Werren 2011; Dalla Benetta et al. 2019a); B chromosomes are non-essential for the host. Yet they are maintained in populations, often in spite of neutral or even deleterious fitness effects, through meiotic drive mechanisms (Jones 1991; Houben 2017). These mechanisms are diverse and with few exceptions remain poorly understood. Many Bs appear to exploit existing asymmetries in meiosis: for example, securing their inclusion into oocyte rather than polar bodies during oogenesis in animals (e.g. in a locust, Pardo et al., 1994) or into the generative nucleus rather than the tube nucleus during pollen formation in plants (e.g. in rye, Banaei-Moghaddam et al., 2012). In other cases, the drive mechanism exploits asymmetries in transmission of chromosomes from parents to offspring: In the haplodiploid wasp *Nasonia vitripennis*, haploid males normally do not pass genetic material on to male offspring. However a sperm-derived B chromosome in this species drives by eliminating all other paternally-inherited autosomes from fertilised female embryos during embryogenesis. This turns these embryos into haploidised males which will incorporate the B chromosome into their sperm (Reed and Werren 1995; Swim et al. 2012; Dalla Benetta et al. 2019b).

However, probably the most extreme example of B chromosomes taking advantage of asymmetries in reproduction is found in a small plant feeding hemipteran insect, the obscure mealybug *Pseudococcus viburni (*Hemiptera: Pseudococcidae). Here the B chromosome hijacks their peculiar reproductive system that involves non-Mendelian chromosome transmission to increase its own transmission: Mealybugs reproduce via paternal genome elimination (PGE), in which maternally-inherited chromosomes drive at the expense of paternally-inherited chromosomes during male meiosis, which are eliminated from sperm (Bongiorni et al., 2004; Brown and Nur, 1964; Dallai, 2012; Nur, 1990). The B chromosome in *P. viburni* drives by avoiding elimination when paternally-derived and is included in all sperm (Nur 1962; Nur 1966; Nur 1966; Nur 1969). So in this species B chromosomes increase their transmission by avoiding meiotic drive (PGE constitutes whole-genome post-meiotic drive), rather than inducing it themselves. As a result, studying the drive mechanisms of this B chromosome not only provides potential insights into the way a single-chromosome reproductive parasite can increase its transmission, but might also help unravel the mechanisms that govern paternal genome elimination, a type of reproduction that is poorly understood despite its widespread distribution and repeated evolution (seven origins and 10,000s species) across Arthropods (Gardner and Ross 2014; de la Filia et al. 2015).

Cytogenetic studies have provided a detailed description of the behaviour of the B chromosome in *P*.*viburni* (summarized in Figure 1), while those in a number of other mealybug species give further insights into the behaviour of maternal and paternal chromosomes. In males of all mealybugs without B chromosomes, under PGE all paternally-inherited chromosomes are eliminated during spermatogenesis (Nur 1980 and citations therein). In addition, in most male tissues paternal chromosomes are heavily condensed through heterochromatinization and are transcriptionally suppressed (Brown and Nur 1964; Bongiorni et al. 2001; de la Filia et al. 2020, Figure 1). During male meiosis, chromosomes segregate based on their parental origin during anaphase II, which involves a monopolar spindle that only interacts with the euchromatic maternal set while the heterochromatic paternal complement lags behind (Bongiorni et al. 2004). Only spermatids containing maternal chromosomes mature into spermatozoa, while those containing heterochromatic paternal chromosomes degrade (Hughes-Schrader 1948; Nur 1962; Bongiorni et al. 2004; Bongiorni et al. 2009). B chromosomes found in populations of *P. viburni* appear to exploit this unusual inheritance pattern as an accumulation mechanism. Regardless of its parental origin, the B shows the same heterochromatic state as the paternal chromosomes in somatic tissues of males (Figure 1&2). However, the B becomes fully decondensed at the beginning of male meiosis, thus mimicking the maternal chromatin state, and migrates with the maternal chromosomes even when inherited from the father (Nur, 1962 and Figure 1&2). By doing so it avoids elimination by always securing inclusion into the two maturing spermatocytes (instead of only when maternally-derived), rather than into the other two degrading nuclei. Via this accumulation mechanism, the B can reach a transmission rate as high as >0.9 through males in spite of clear fitness consequences, such as delayed developmental time and sperm depletion (Nur, 1969; Nur, 1966; Nur, 1966). The B chromosome is also transmitted through females although it does not show any drive, in fact transmission through female meiosis is slightly <0.5 (Nur, 1962, Figure 1). The ability of the paternal B chromosome to increase its transmission in males also varies among different “host” genotypes and studies have shown that it is the maternal rather than the paternal genome that determines this variability in transmission rate (Nur and Brett 1985; Nur and Brett 1988).

**Figure 1.**
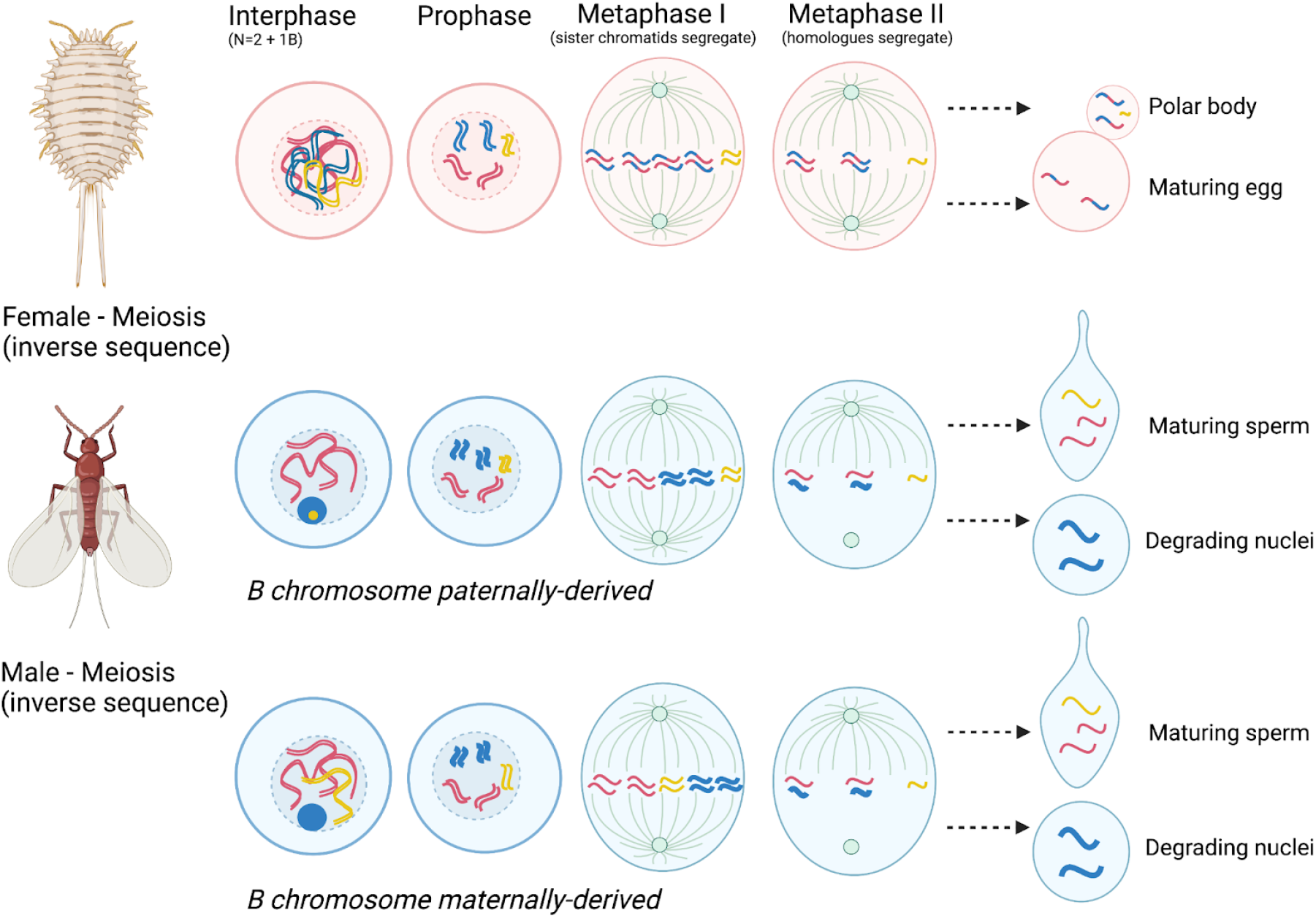
A schematic depiction of the behaviour of the B chromosome during male and female meiosis and gametogenesis in *Pseudococcus viburni*. The top panel shows female meiosis, the bottom two panels show male meiosis with two scenarios where the B is either paternally-derived (top) or maternally-derived (bottom). Throughout the chromosome sequence is shown for N=2 autosomes (maternally-derived autosomes in pink, paternally-derived autosomes in blue) and one unpaired B chromosome (in yellow). Please note that mealybugs have an inverted meiotic sequence and have holocentric chromosomes without a localized centromere. During interhase, prophase each chromosome consists of two loosely connected chromatids that segregate at meiosis I into two daughter cells. For simplicity only one of these daughter cells is depicted in meiosis II and as a result only two rather than four gametes are shown. In male meiosis paternally-derived chromosomes form a condensed heterogametic body in interphase, but separate at prophase and during the two meiotic divisions, however throughout the entire sequence paternal chromosomes appear more condensed than maternal chromosomes. Maternally-derived B chromosomes behave like the other maternally-derived chromosomes and are always included into the maturing sperm nuclei. Paternally-derived Bs are less condensed than the paternal A chromosomes during meiosis and tend to segregate with the less condensed maternal chromosomes into the maturing sperm nuclei, rather than with the paternally-derived chromosomes which segregate into the degrading nuclei.

Based on these cytogenetic studies we hypothesize that the B chromosomes in *P*.*viburni* escape elimination by changing their chromatin structure thereby enhancing their transmission. However up until now we do not have any genetic data that might help us test this hypothesis and help to better understand the molecular underpinnings of the Bs drive. Here we redress this lack of data by sequencing and comparing genomes of individuals with and without B chromosomes. We combine these data with a large gene expression study comparing the expression of B-linked genes during male and female meiosis and gametogenesis. We expect genes involved in drive to be upregulated in males as the B does not show drive in females. The few similar studies in other species with B chromosomes to date show that most B chromosomes do contain actively transcribed genes, but are often gene-poor relative to autosomes and highly enriched for repetitive sequences (as reviewed by Ahmad and Martins 2019). In a few cases, these studies were actually able to identify B-linked genes responsible for the drive, such as the gene *haplodizer* in the wasp *N. vitripennis* (Dalla Benetta et al. 2020). We present a highly contiguous *de novo* draft genome for an inbred *P. viburni* line which contains a single B chromosome. By comparing this reference genome to genome data from lines without Bs and with 2 B copies we were able to identify and assemble 1.37Mb of B-linked sequence containing 78 protein coding genes. We found that while most of these genes showed low levels of expression, we identified a number of promising B-linked candidate genes which have annotated functions in meiosis and histone modification and are expressed during gametogenesis. Most notable is the histone acetyltransferase Rtt109/CBP which has a putative role in meiosis and is upregulated in males compared to females. Finally, we investigate the relationship of the B chromosome to the rest of the core genome and find that autosomal paralogs of B-linked genes are equally distributed across the genome. This suggests that the B is not derived from one particular autosome.

## Results

### *B chromosomes in* P. viburni *populations*

We collected *P. viburni* from glasshouses at the Royal Botanical Gardens in Edinburgh, Scotlan and established isofemale lines. To identify B- and B+ *P. viburni* lines for DNA and RNA-seq (Supplementary Table S1), we stained and counted chromosomes in embryos. We found that one of our *P. viburni* populations, RBGE25, did not carry a B chromosome (Fig. 1A(i)) and established two B- experimental lines: PV21 and PV23. We then selected two other populations that carried either one B, RBGE31 (Fig. 1A(ii); from which we established an experimental line, PV13) or on average two Bs, RBGE29 (Fig. 1A(iii), experimental line PV04).

### B chromosome behaviour during meiosis

We used cytological preparations of second-instar male dissected testes to observe how the B chromosome behaves during meiosis and spermiogenesis and compare this to the earlier studies which were done in a different population. At the beginning of meiosis, the maternal genome (MG) starts condensing to form metaphase chromosomes, but since the paternal genome (PG) is already in the heterochromatin state, the paternal chromosomes start to individualize before distinct maternal chromosomes become apparent (Fig. 1B(i)). In B-populations, we observed n=5 paternal chromosomes. In B+ populations, during the same early meiosis stage we could observe n=5+1/2Bs paternal chromosome in the heterochromatic side of the nucleus (Fig. 1B(iii) carrying 2Bs). Male meiosis in mealybugs results in a quadrinucleate cell (four nuclei within a shared cytoplasm). This cell contains two opposite nuclei containing the heterochromatic PG and two opposite nuclei with the euchromatic MG. In B-populations, we could clearly see the n=5 PG chromosomes (Fig. 1B(ii)), while in the B+ populations, the B chromosome(s) either segregates with the MG or the PG (Fig. 1B(iv, v)). However in both 1B and 2B populations we saw that generally the nuclei containing the MG carried 1-2 Bs at the end of meiosis, observed as condensed dots within the nuclei (not shown, but see 1B(v)), while segregation with the PG genome was rare. After meiosis the quadrinucleate cell elongates, resulting in the elongation of the MG nuclei only (Fig. 1B (vi and vii)). The PG remains condensed and is eventually eliminated. So it appears that even paternally-derived B chromosomes that colocalize with the PG at the beginning of meiosis segregate with the MG by the beginning of spermiogenesis, when the PG is eliminated. As such, B chromosomes escape PGE despite their paternal origin.

### Genome assembly

We generated a *P. viburni* genome assembly from pooled females of the B-carrying experimental line PV13. We assembled 28.6Gb of PacBio subreads and used two Illumina short read libraries generated from three females from the same line to polish the assembly (see Materials and Methods and Supplementary Table S1 for a summary of our sequencing efforts). Our genome assembly, after removing contigs derived from the bacterial endosymbionts housed by the species (International Aphid Genomics Consortium 2010; Chen et al. 2016), comprises 435.6Mb in 2,392 scaffolds with a GC content of 33.7% and a N50 of ∼0.87Mb. These and other assembly metrics are shown in Table 1.

**Table 1.**
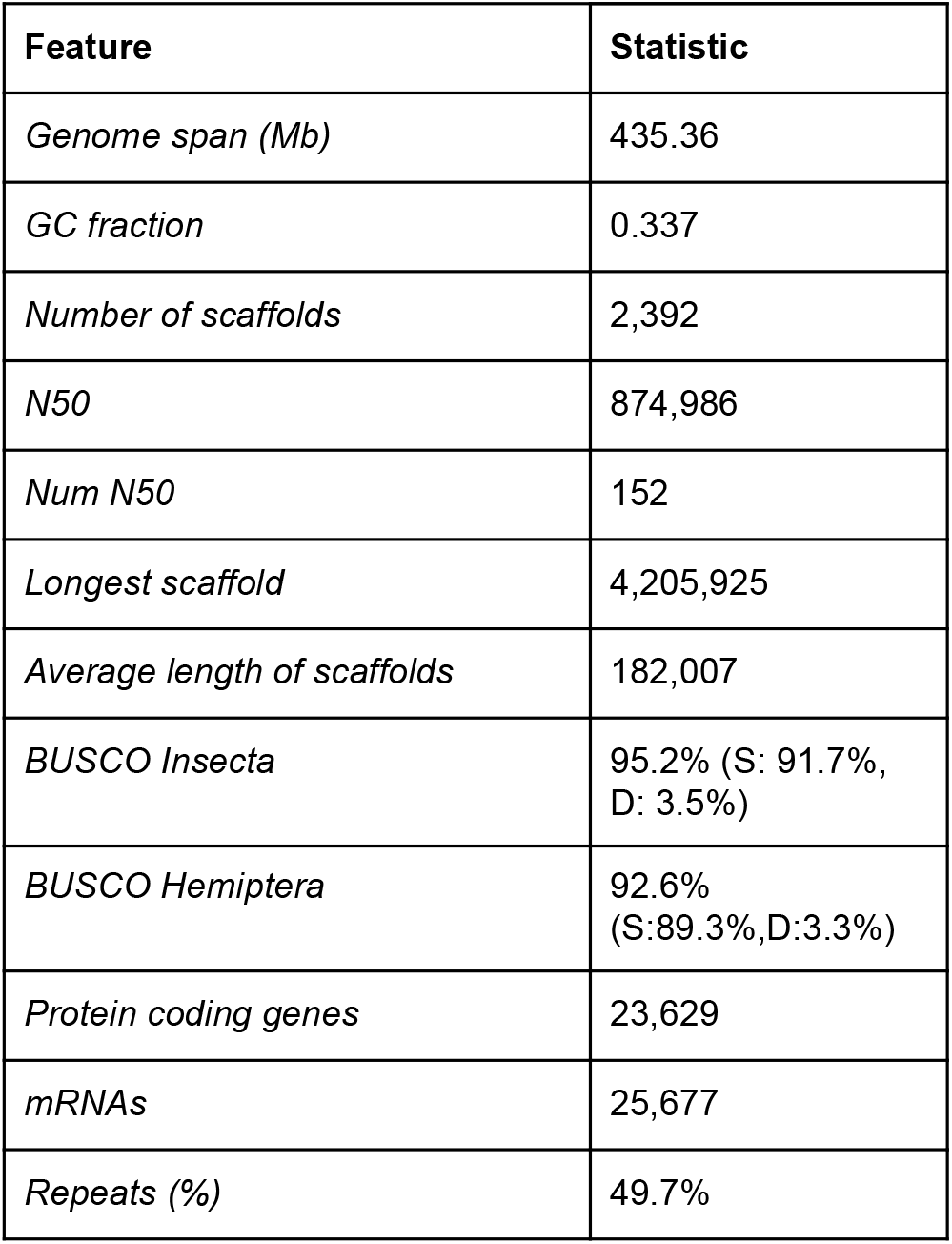
Genome assembly and annotation statistics

Our genome assembly shows high completeness, as revealed by the presence of >90% single copy orthologs from the insect and hemipteran BUSCO gene sets. Half of the genome (216.2Mb) consists of repetitive sequences, higher than the 38-45% range estimated in *Phenacoccus solenopsis* (Li et al., 2020) and *Pseudococcus longispinus* (Garber et al. 2021 publicly available at https://ensembl.mealybug.org/Pseudococcus_longispinus_v1)) and hemipterans such as *Acyrthosiphon pisum* or *Bemisia tabaci* (International Aphid Genomics Consortium, 2010; Chen et al., 2016). The vast majority of repeats (96.6%) are unclassified. We predicted a set of 23,629 protein coding genes, which is twice the number found in the most complete mealybug genome available to date, *P. solenopsis*, but very similar to the gene estimate in the more closely related *P. longispinus*. We annotated 13,515 genes using homology searches and functional domain prediction. The BUSCO scores for our annotated gene set were 95.1% (Insecta) and 93.1% (Hemiptera), which were extremely close to our assembly benchmarking and therefore indicative of a successful gene annotation. Moreover, less than 5% of BUSCOs were duplicated, suggesting our higher gene count is probably not the result of systematic assembly errors.

### Identifying B-derived scaffolds in the reference genome

We reasoned that B-derived Illumina reads might either be unique to B chromosomes or shared between A and B chromosomes, as is common in supernumerary chromosomes (Ahmad and Martins 2019). Therefore, we combined several lines of evidence to assign scaffolds to autosomal (A) and B chromosomes using the short read libraries generated for the two B+ (PV13 and PV04) and B-lines (PV21 and PV23) (see Materials and Methods for full details).

First, we mapped the short reads to our reference assembly and identified scaffolds with higher coverage in B+ lines compared to B-lines using two coverage thresholds: a strict threshold of log2(coverage ratio) ≥ 2 (145 scaffolds) and a more relaxed threshold of log2 ≥ 0.58, corresponding to a 3:2 ratio that could be expected for scaffolds shared between autosomes and a single copy of the B chromosome (a further 136 scaffolds) (Fig. 2A, Supplementary Figure S1). Second, we assembled the short reads for each library into reference-independent assemblies and aligned them to our reference assembly to identify scaffolds in the reference genome present in the PV13 and PV04 Illumina assemblies but not in PV21 or PV23 (107 scaffolds) (Fig. 2B, Supplementary Table S2). Third, we used a kmer-based approach to identify overrepresented kmers in the B+ libraries using the 3:2 rule. We recovered 553,816,347 27mers with at least 3 counts in all libraries, of which 49.78M (∼9%) were assigned as putative B kmers. We mapped both sets of B and non-B kmers to the PacBio assembly and identified 58 scaffolds with at least as many putative B as non-B kmer mappings (Fig. 2C). We further reasoned that B-linked sequences should be present in higher coverage in PV04 (the 2B line) than in PV13 (the 1B line), although we did not incorporate this as a formalised rule in our decision process since B number can vary within lines and we did not directly karyotype the sequenced individuals.

**Figure 2.**
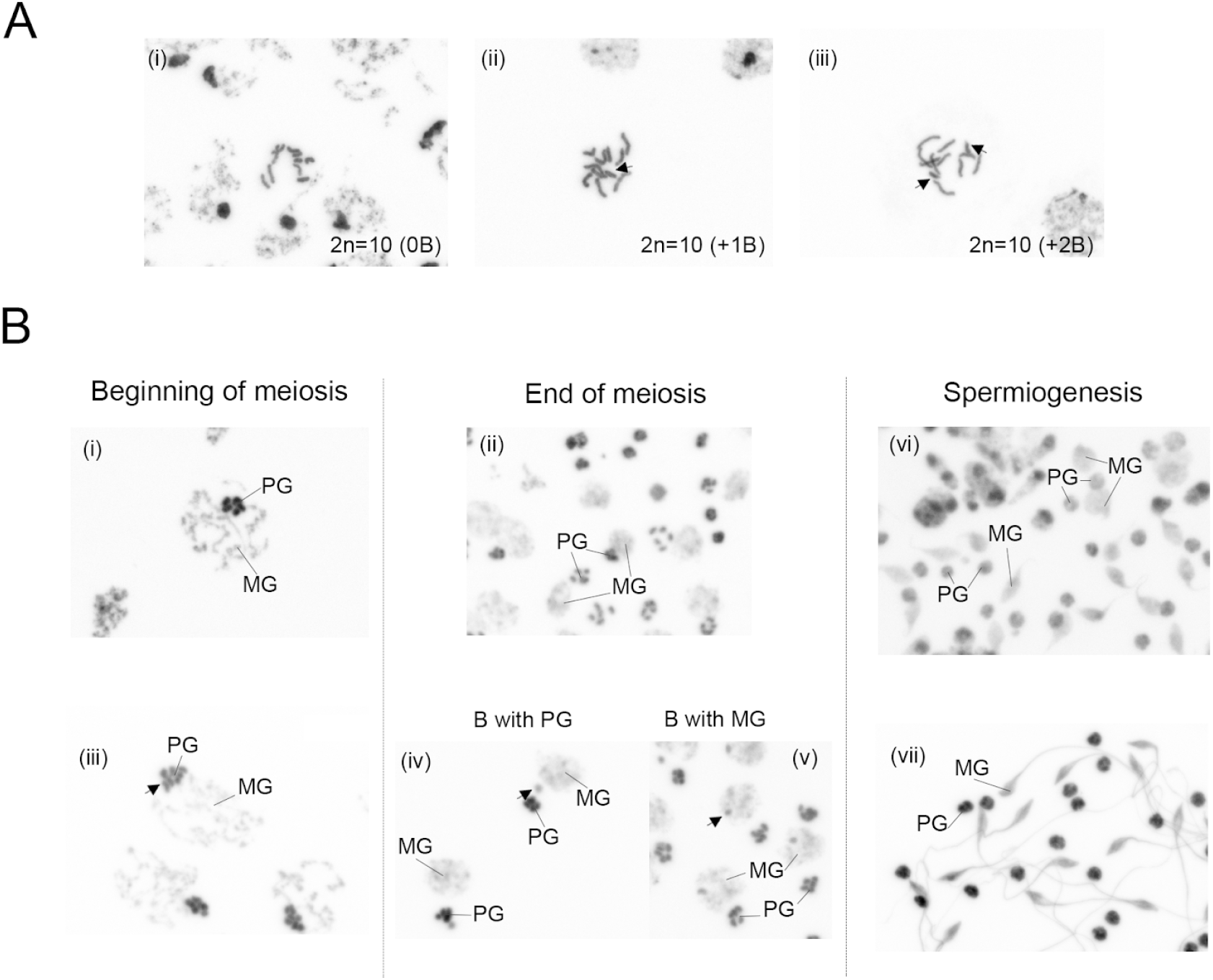
Cytological preparations of screened *Pseudococcus viburni* populations used for sequencing. (A) Embryonic cells with (i) only A chromosomes 2n=10, from PV23. (ii) A chromosomes 2n=10 + 1B chromosome, from PV13, (iii) A chromosomes 2n=10 + 2B chromosomes, from PV04. (B) Testes cells of second-instar males with cells at the beginning of meiosis, where the paternal chromosomes are showing either n=5 from PV23 (i) or n=5+2B from PV04 (ii); and cells at the end of meiosis, where the future gametes do not carry B in 0B lines (PV23) form (iii) or where the B chromosome either segregates with the PG (iv) or the MG (v) from PV13. During spermiogenesis, the PG becomes heterochromatinised again, while the MG decondenses and the nucleus starts elongation to form the future gametes (vi and vii), from PV13. All images are at the same scale (x40 or x20), and only show DNA, cells stained with DAPI

Combining these different criteria, we were able to assign B status to reference scaffolds with varying degrees of confidence (B1 = high confidence, B2 and B3 = lower confidence), with the highest confidence set, B1, comprising 103 scaffolds and 1.37Mb of reference sequence (Fig. 2D, Table 2, Supplementary File S1.0). Approximately half of the B1 scaffolds are larger than 10Kb, with the largest scaffold spanning 190Kb (Supplementary Figure S2).

**Table 2.** *Assignments of A and B scaffolds in the* P. viburni *reference assembly*.

| Set* | N scaffolds | Mb | Mean PV04/PV13 cov ratio | Predicted genes (of which annotated) |
| --- | --- | --- | --- | --- |
| A | 2084 | 428.2 | 1.01 (0.64 SD) | 23143 (13309, 58%) |
| B set 1 | 103 | 1.37 | 2.99 (0.84 SD) | 78 (32, 41%) |
| B set 2 | 106 | 3.16 | 1.81 (0.90 SD) | 297 (112, 37%) |
| B set 3 | 99 | 2.58 | 0.51 (0.28 SD) | 111 (62, 56%) |

\* Assignment strategy as follows: **B set 1 (highest confidence)**, log2 read coverage ratio between all B+/B- pairs $\geq 2$ and coverage in PV04 > coverage in PV13 and 1-1 mapping to a contig in the B+ Illumina assemblies only or log2 putative A/B kmer ratio $\geq 0$ . **B set 2**, cov in PV04 > cov in PV13 and at least one of the following: log2 cov ratio between all B+/B- pairs $\geq 0.58$ , 1-1 mapping in B+ assemblies only, log2 A/B kmer ratio $\geq 0$ . **B set 3 (lowest confidence)**, at least one of the following: log2 cov ratio between all B+/B- pairs $\geq 0.58$ , 1-1 mapping in B+ assemblies only, log2 A/B kmer ratio $\geq 0$ .

We used PCR to validate these results by targeting the largest B1 scaffold, scaffold_552. We designed three sets of primers (Supplementary Table S3) and genotyped females from the four experimental lines and another mealybug species, *Planococcus citri*, as a negative control. As expected, only PV13 and PV04 individuals showed amplification (Supplementary Figure S7).

### Composition of B-linked sequences

B chromosomes are typically rich in repetitive sequences. Accordingly, we found a higher repetitive component in the highest confidence B1 scaffolds, 71.9%, compared to the whole genome (see S1.15 and 1.16 in the supplement file), with the vast majority of these repeats unclassified. However it is likely that this substantially underestimates the true repeat content of the B as the assembled scaffolds might not capture highly repetitive regions. Therefore we also analysed the repeat content of the raw reads from the B- and B+ libraries, following a pipeline in Ruiz-Ruano et al. (2016). Across all libraries we identified 19 satellite repeats (Figure 4, Repeat Analysis Supplement), four of which (with repeat-length ranging from 30-414) show a strong over-abundance in B+ libraries. One 172bp B-linked repeat is particularly abundant, comprising 3.9% of all reads in the 2B library and 1.3% in the 1B library. Divergence patterns between copies of this repeat suggest that much of its abundance is due to recent extensive amplification (Figure 4).

We identified 78 protein-coding genes (81 transcripts) located on 26 of the 103 B1 scaffolds (Supplementary file S1.1). Most of the genes are located on the largest scaffold (scaffold_552). Additionally, 408 genes are located on the lower-confidence B2 and B3 scaffolds, of which 174 are annotated (Supplementary file S1.3). We tested whether the B chromosome shows a clear homology to the rest of the genome using reciprocal blast of all the B-linked scaffolds against the core genes annotated in the assembly, and furthermore against all annotated genes of closely related species *P. solenopsis* (Li et al. 2020) and *P. longispinus* (Garber et al. 2021). Of all 486 B-linked genes we identified 56 that carry a paralog in the core genome (Supplementary Table S6). Furthermore for 43 and 11 B-linked genes we identified orthologs in *P. longispinus* and *P. solenopsis* respectively (Supplementary Table S6). In the case of *P. solenopsis* we were able to identify the orthologs located on all 5 chromosomes. The low numbers of identified paralogs and orthologs, together with scattered hits across the *P. solenopsis* suggest the B chromosome is not derived from a core genome chromosome. Additionally, we found no support that the B chromosome is homologous to a *Phenococcus* core chromosome using nucleotide-alignment to *P. solenopsis* genome (Li et al. 2020). We found none of the scaffolds in the most stringent putative B chromosome set (B1) showed conserved homology with any of the five chromosomes. Even for relaxed criteria (B2, B3) there was almost no detectable synteny (Supplementary Table S6). This is unlikely a result of methodological approach as we were able to find homologous chromosomes of 80% of the *P. viburni* core genome scaffolds (see Supplementary Table S6). To test for a more recent origin, we also compared *P. viburni* to the more closely related congener, *Pseudococcus longispinus*. More putative B linked scaffolds have syntenic alignments with *P. longispinus* than with *P. solenopsis*, but even between the congeners there is little evidence of long tracts of conserved sequence between the *P. viburni* B and the A chromosomes of other mealybug species (Supplementary Table S7, Figure S5).

Furthermore, for 17 B-linked genes we identified a homolog in at least two out of three core genomes. In all 17 cases these genes showed patterns consistent with intraspecific origins. Divergences in all cases followed phylogenetic relationships of the three species - the highest similarity to *P. viburni* core genome, followed by similarity to *P. longispinus* and lowest similarity to *P. solenopsis*. Although this pattern of nucleotide similarity is consistent with intraspecific origin, we found a homologous sequence only for a relatively small fraction of B-linked genes - 93 out of a total 486 predicted genes (those in the B1, B2 and B3 criteria). To further investigate possible other sources we searched for homologies against NR, NCBI, and UniProt databases using the same strategy used by blobtools2 (https://github.com/blobtoolkit/blobtools2; Laetsch and Blaxter, 2017). We found a significant hit for additional 104 out of 394 genes of unknown origin. Majority of homologs were found in Arthropods, and most of them specifically in Hemiptera (Supplementary Table S5). One notable exception is scaffold_360, which carries a set of 18 viral genes as well as six arthropod genes and two paralogs to the core genome.

### Putative functions of B-linked genes

In our genome annotation, thirty-two B1 genes appear annotated either with protein family domains and/or significant homology to protein databases. Among the functions associated with these genes, we found processes such as microtubule motor activity, ion transport, transmembrane transport, histone modifications, transcription regulation, DNA integration and transposition activity (Supplementary file S1.1). Additionally, 408 genes are located on the lower-confidence B2 and B3 scaffolds, of which 174 are annotated (Supplementary file S1.3).

Our RNA-seq data of males and females from the four experimental lines revealed, however, that most B1 genes are not expressed in second instars. Only 13 B1 genes are expressed in at least one of the sample groups (TPM>1), of which three have expression levels about 10 TPM (Supplementary file S1.4). When averaging across all B+ and B-samples, we found that B1 genes show significantly lower expression than B2 and B3 genes in both cases (Supplementary Figure S3).

### Sex-biased gene expression

Before exploring whether the presence of B chromosomes can differentially affect males and females, particularly during meiosis, we compared the gene expression profiles of our male and female RNA-seq samples in order to identify sex-biased expression and confirm we captured the right developmental stage. The 2nd instar stage is critical as it corresponds to the onset of reproductive tissue maturation, but also to the establishment of diverging developmental pathways: at later larval stages and adulthood, mealybugs exhibit striking sexual dimorphism (Vea et al. 2016; Bain et al. 2021)). Therefore, our RNA-seq data gives us the opportunity to directly examine sex-specific differences in gene expression at late 2nd instar stage (Fig. 3A).

**Figure 3.**
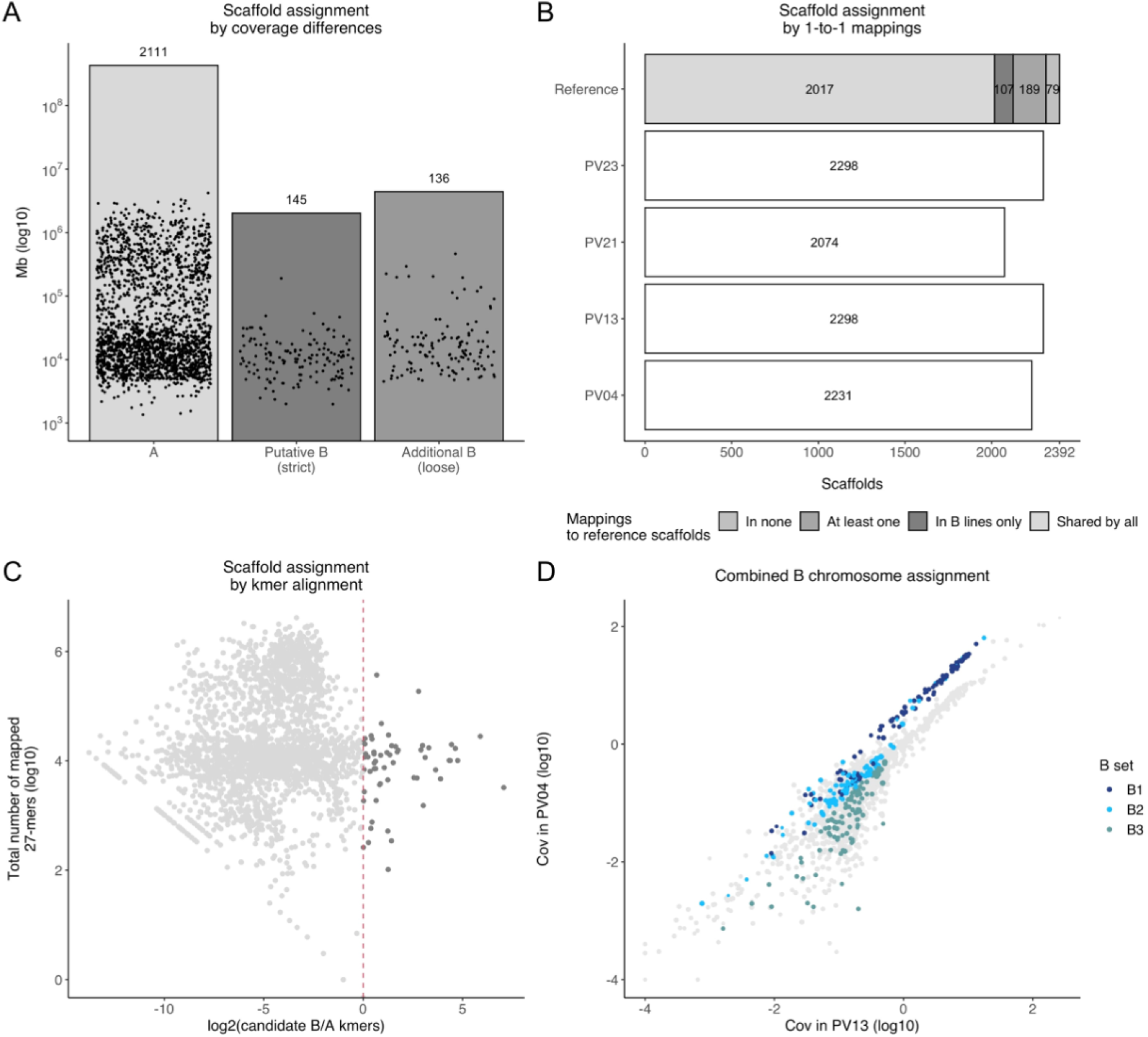
B-linked scaffold assignment. (A) Assignment based on read coverage differences between B+ and B-lines. (B) Assignment based on mappings to individual Illumina assemblies of the individual lines. Counts of scaffolds with 1-to-1 mappings are shown on the X axis. (C) Assignment based on alignment of kmers overpressented in B+ lines. (D) Scatterplot of scaffold read coverage in the two B+ lines, PV04 (2B) and PV13 (1B). B-assigned scaffolds are represented in shades of blue according to assignment confidence. A chromosomes are shown in light gray.

**Figure 4.**
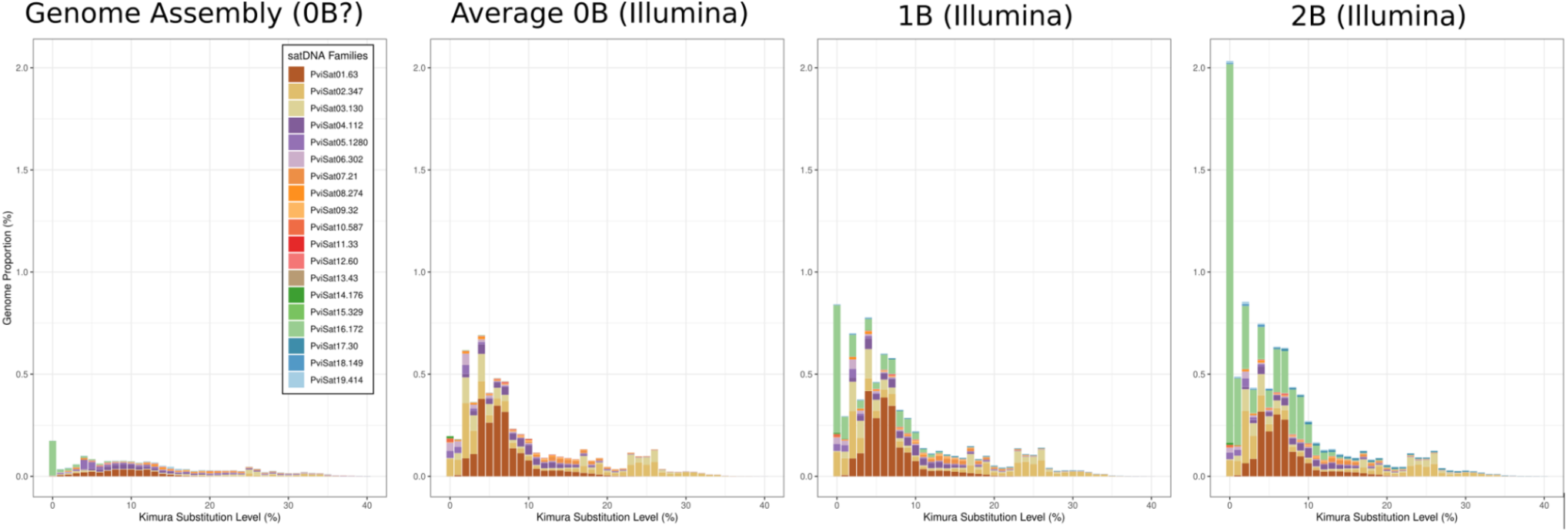
Satellite repeats. Repeat landscapes for the assembly and each separate library (average across two 0Bs, 1B and 2B). Each plot shows the abundance for each of the 19 satellite repeats at different divergence intervals (Kimura substitution levels). The satellites are numbered in decreasing order of 0B abundance and with the monomer size at the end of the name. The 1B and 2B libraries are enriched for the PviSat05-1280 and in particular the PviSat16-172 satellites. PviSat16-172 shows very low divergence between repeats indicating recent amplification.

**Fig 5.**
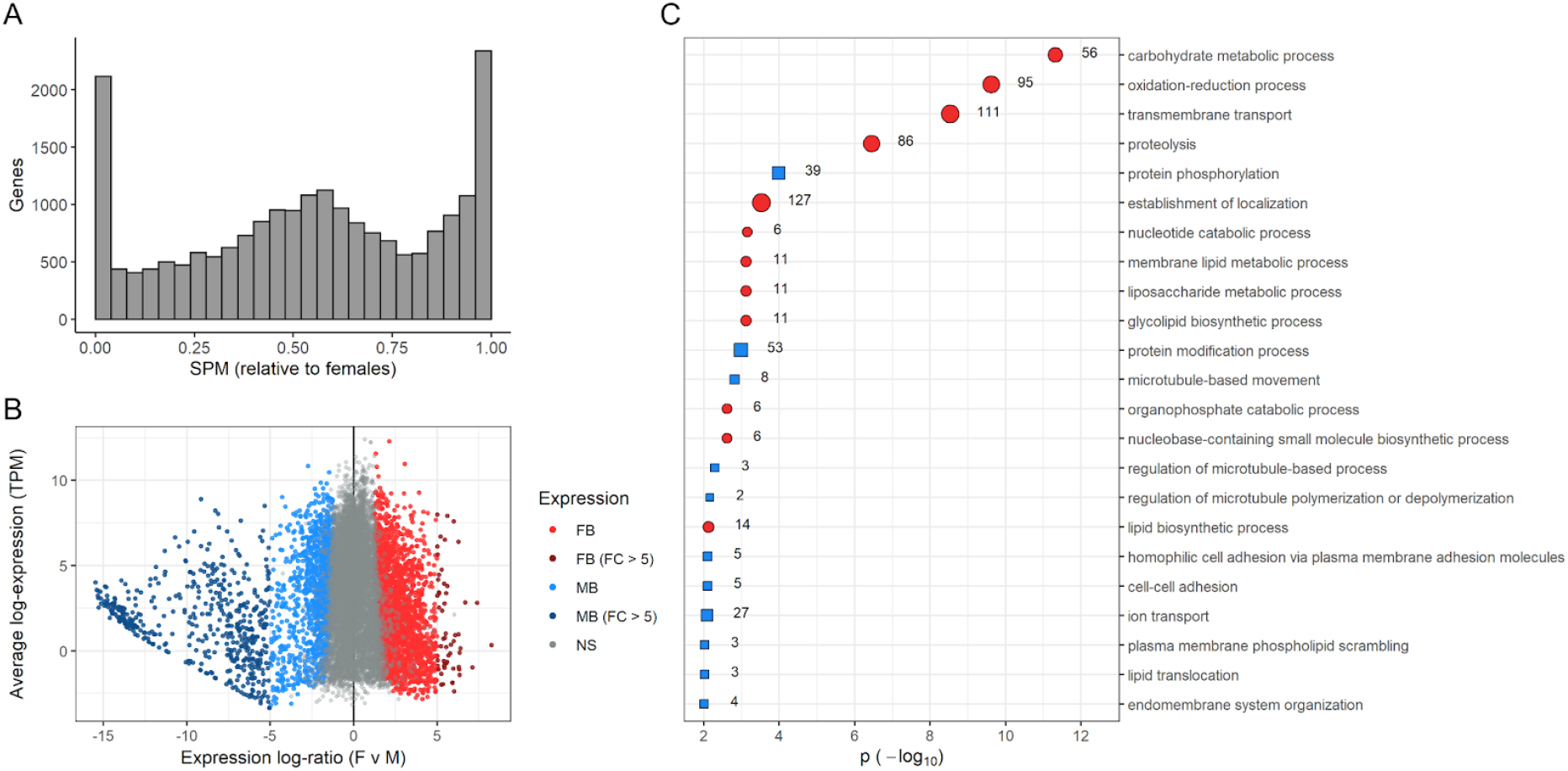
Sex-biased expression in 2nd instars of *P. viburni*. (A) Histogram of sex-specificity (SPM, a proportion of expression ranging from 0 (in males only) to 1 (in females only) (Xiao et al. 2010; Kryuchkova-Mostacci and Robinson-Rechavi 2017). (B) Scatterplot of expression fold-changes in genes and average levels of expression. Significantly differentially expressed genes are shown in red (female-biased) and blue (male-biased). Strongly sex-biased genes (|FC >5|) are shown in darker shades. (C) Significantly enriched GO terms related to biological processes in females (red circles) and males (blue squares). 803 female-biased genes and 653 male-biased genes had GO terms associated with them. The background population consisted of 6,968 genes included in the differential expression analysis with GO terms.

**Fig 6.**
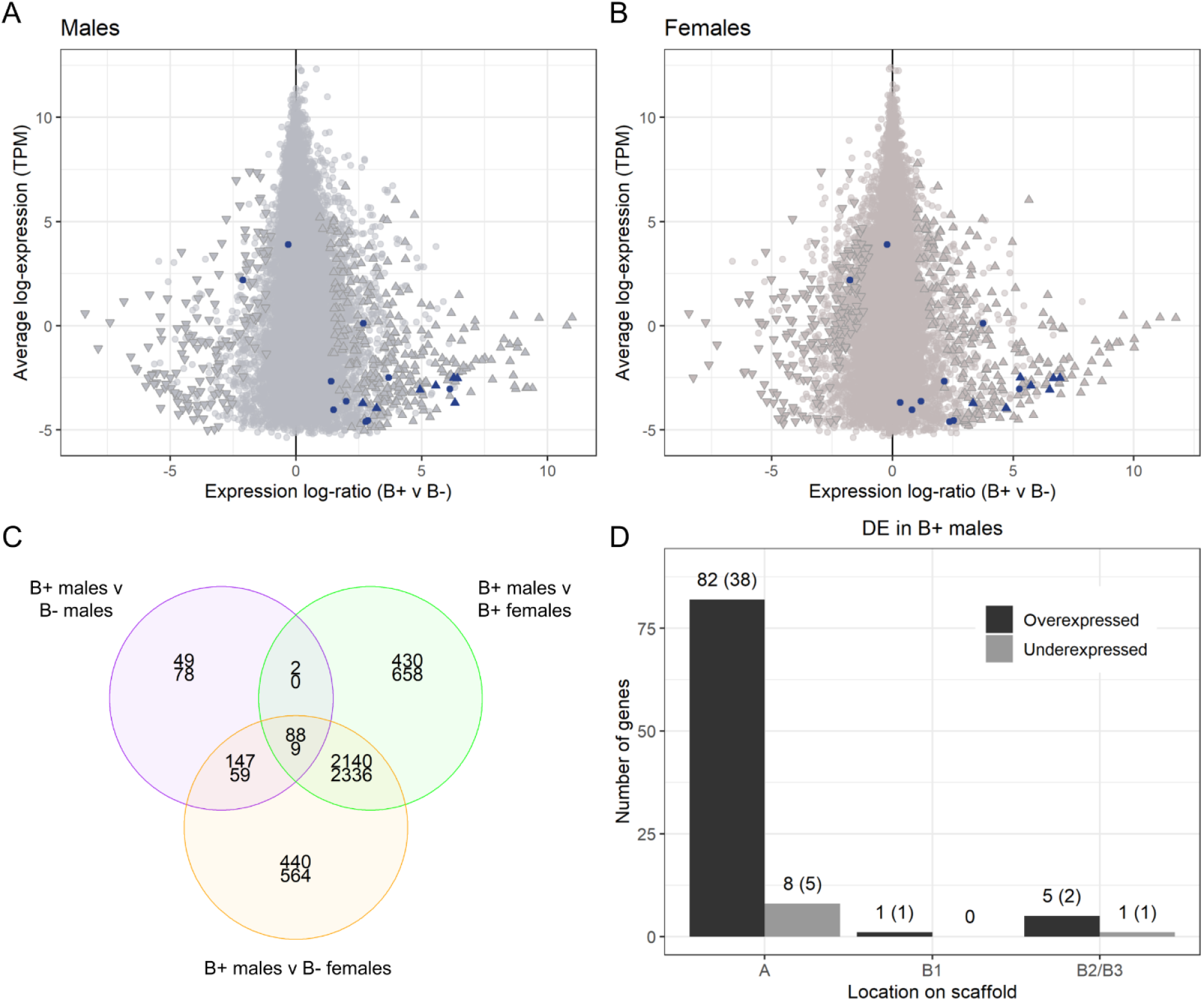
Differential gene expression analysis depending on presence or absence of B chromosomes in 2nd instar mealybugs. (A) Scatterplot of expression fold-changes in genes and average levels of expression between B+ and B-males. Upwards and downwards arrowheads represent genes that are significantly overexpressed and underexpressed, respectively, in B+ males. Genes located on B1 scaffolds are shown in dark blue. (B) Scatterplot of expression fold-changes in genes and average levels of expression between B+ and B-females. (C) Venn diagram showing overlaps in differentially expressed genes between B+ males and the other three groups: B-males, B+ females and B-females. Counts of overexpressed genes in B+ males shown above; underexpressed genes below. (D) Number of differentially expressed genes between B+ males and the other three sample groups grouped by location on the genome assembly. Number of annotated genes in each group are also shown in brackets.

Thus, we conducted a differential expression analysis between all male and female RNA-seq samples in our study. Out of a total of 15,673 genes with enough expression in at least one sex to be included in the analysis (Materials and methods), we identified 3,830 genes (24.4%) with sex-biased expression (FDR < 0.05, |logFC > 1|) (Fig. 3B, Supplementary file S1.5). Of these, 2,186 are overexpressed in females and 1,644 in males, of which 57 and 484, respectively, can be considered strongly sex-biased (logFC > 5). We conducted a GO enrichment analysis of sex-biased genes in both sexes (p < 0.01, Fig. 3C) and found that female-biased genes are enriched in metabolic processes such as lipid and carbohydrate metabolism (Supplementary file S1.6), while male-biased genes are enriched in protein modification, membrane organisation, lipid translocation and microtubule organisation (Supplementary file S1.7).

Among the male-biased genes, a few are involved in endocrine pathways linked to insect metamorphosis. Particularly, g2298 codes for JHAMT protein, the enzyme involved in the last step of juvenile hormone (JH) synthesis. This transcript was previously found to have diverging expression profiles between males and females at the onset of metamorphosis in mealybugs (Vea et al. 2016). However, we only found a single strongly male-biased gene that is directly related to reproduction: g17352 (protamine-like sperm chromatin condensation).

### Differential gene expression between B-carrying and non B samples

We then examined differential expression between B-carrying and B-sample groups with a focus on B+ males, due to the distinct behaviour and meiotic drive of the B chromosome in males. To do so, we performed a differential expression analysis for 18,066 genes which included four contrasts: B-carrying males versus the other three sample groups (B-males, B+ females and B-females) and, additionally, B+ females versus B-females (18,066 genes with significantly differential expression set to |log2FC > 0.58| and FDR < 0.01; Table 3; Supplementary Figure S4; Supplementary file S1.8).

**Table 3.**
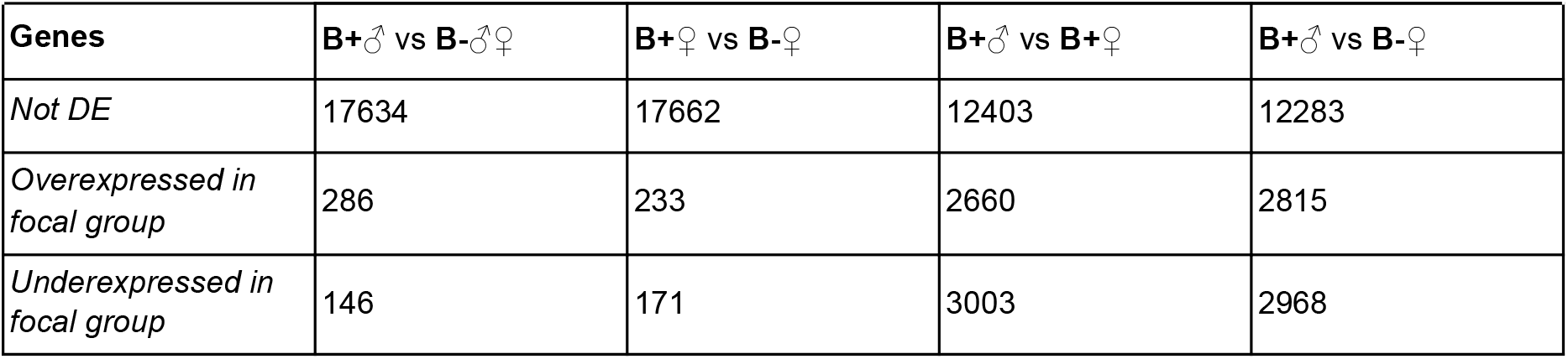
Differentially expressed genes between contrasts (FDR < 0.01, log2FC > 0.58).

We found 432 differentially expressed genes between B+ males and B-males, 232 of which are annotated (Fig 4A; Supplementary file S1.9). Among the 286 genes that are overexpressed in B+ males, we found seven located on B1 scaffolds. These genes include five annotated genes: a histone acetyltransferase Rtt109/CBP (g13953), two fatty acid synthases (g1208, g5582) and two contiguous genes with predicted AAA-ATPase-like domains (g9061, g9062). As expected, we did not find any B1-linked genes among the 146 genes that are overexpressed in B-samples. A GO term enrichment analysis revealed that the differentially expressed genes between B+ and B-males are enriched in four biological processes: DNA integration (GO:0015074), transposition, DNA-mediated (GO:0006313), proteasome assembly (GO:0043248) and nuclear-transcribed mRNA catabolic process, nonsense-mediated decay (GO:0000184) (122 genes with associated GO terms out of 7,375 annotated genes in the background population, Supplementary file S1.10).

The results in females were similar, with 233 genes overexpressed in B+ (7 on B1 scaffolds, including g1208, g5582, g9061 and g9062) and 171 in B-(none on B1) (Fig 4B; Supplementary file S1.11). The differentially expressed genes are enriched in DNA metabolic process (GO:0006259) and telomere maintenance (GO:0000723) (119 genes with associated GO terms out of 7,375 annotated genes in the background population, Supplementary file S1.12). Overall, we found substantial overlap in the annotated differentially expressed genes between B+ males and B-males and B+ females and B-females: 94 genes are upregulated in B+ males and females (Supplementary file S1.13), while 34 genes are downregulated compared to B-males and females (Supplementary file S1.14). We did not find any annotated genes with opposite differential expression patterns in males and females: i.e. upregulated in B+ males and downregulated in B-females, or vice versa.

We then focused on the genes that are differentially expressed in B+ males compared to the other three sample groups (Figs. 4C-D, Table 4). Among the 88 genes that are overexpressed in B+ males, we found a single B1 gene: the histone acetyltransferase g13953. Additionally, we found two annotated genes located on low-confidence B-linked scaffolds: g5312 (B2), a transposase IS4; and g5232 (B3), which contains a N-terminal domain of TAR DNA-Binding Protein 43. The remaining annotated genes are located on A scaffolds and are therefore autosomal. Nine genes are underexpressed in B+ males, including five genes on A scaffolds and a catalase gene, g5248, located on a B3 scaffold.

**Table 4.**
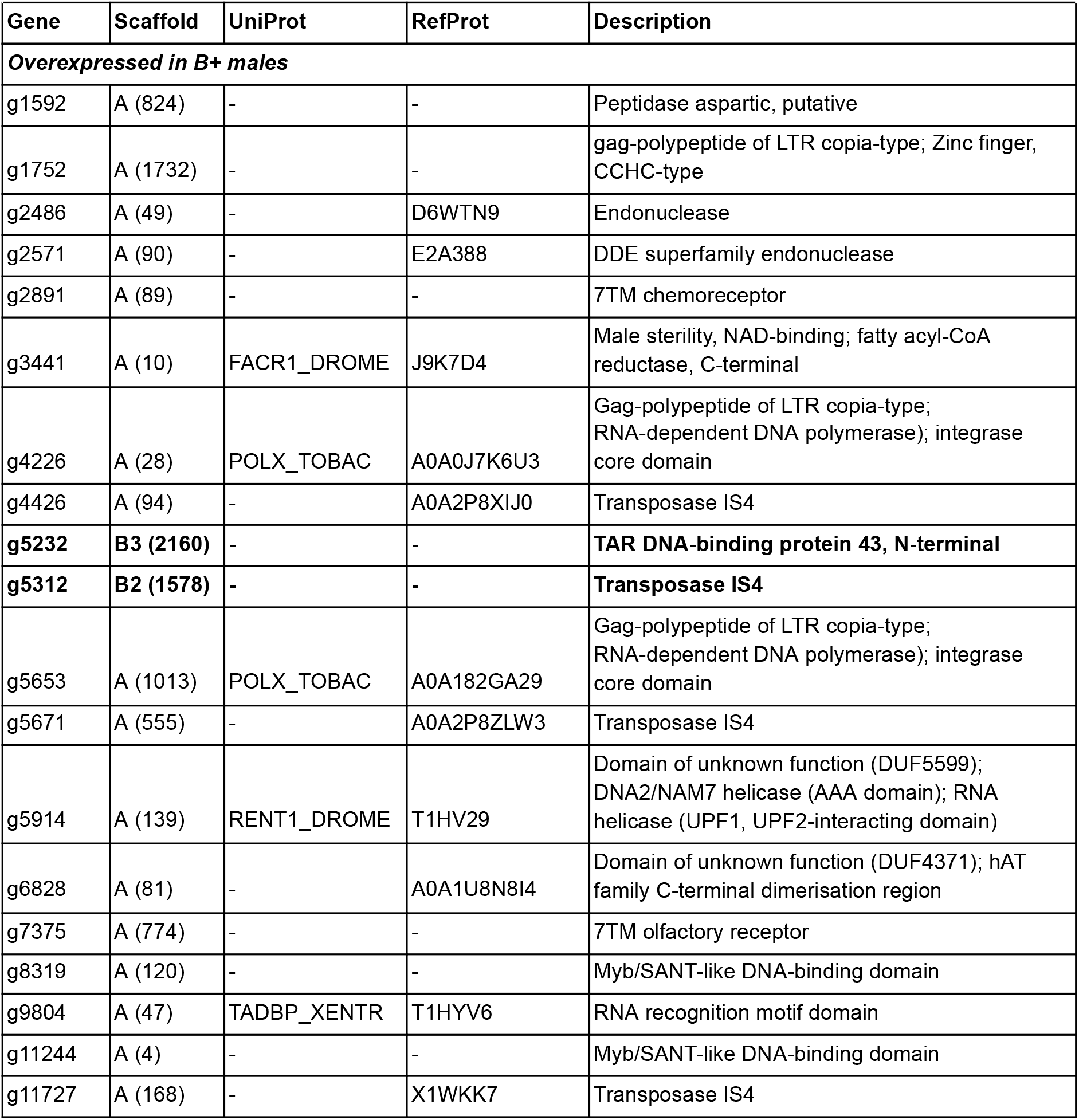
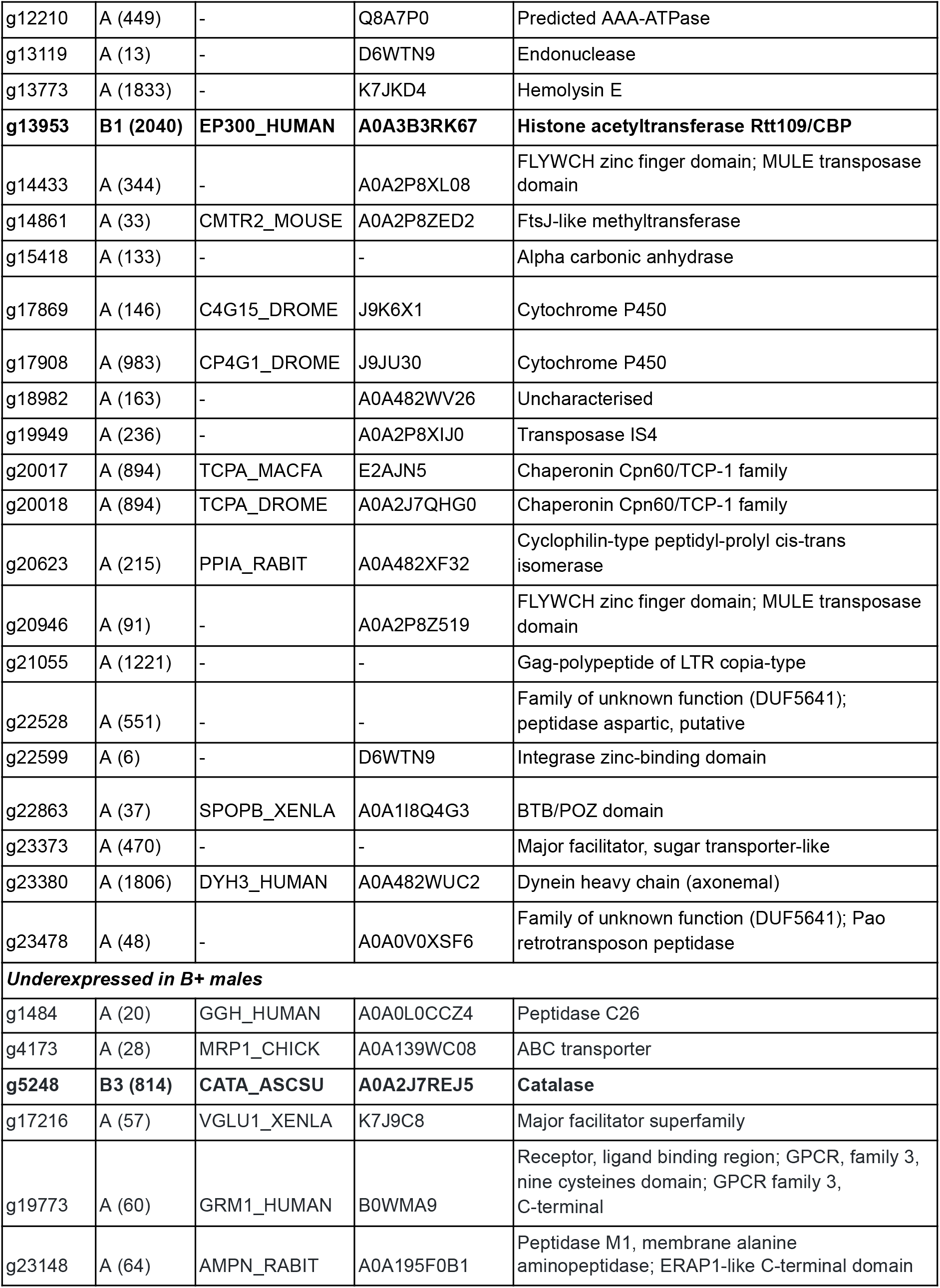
Annotated differentially expressed genes in B-carrying males with respect to the other three groups (B-males, B+ females and B-females), with best BLAST/DIAMOND hits (E < 1e-25) to UniProtKB/SwissProt and reference proteome databases. Genes located on candidate B scaffolds are highlighted in bold. Descriptions are inferred from Pfam/InterPro domains.

In conclusion, the presence of B chromosomes triggers similar transcriptional changes in both males and females, both originating from the B chromosome itself and autosomal genes. Interestingly, the single B gene that is uniquely overexpressed in B+ males is a histone acetyltransferase involved in H3K56Ac, which has a role in meiosis.

### B chromosome transmission and prevalence in two greenhouse populations

The PCR primers that we designed to validate the B status of scaffold_552 also allowed us to easily detect B chromosome presence within individuals. We used primer sets 1 and 2 to survey two glasshouse populations in Edinburgh for the presence, frequency and transmission rate of the B chromosome. The first population at the Royal Botanical Gardens Edinburgh (RBGE), where we sampled two adjacent glasshouses, was the same population where we first discovered the B chromosome and from which we collected specimens that founded the laboratory populations used for this study. The second is an unrelated population at Butterfly World Edinburgh. We identified the prevalence and transmission in each population by taking females into the lab and genotyping 6-14 of their offspring. We found that the B chromosome was present in all glasshouses in both populations, but not fixed: we detected B chromosomes in at least one offspring of all females collected from the first RBGE glass house, but only in 3 out of 5 in the other RBGE glasshouse and in 13 out of 17 in the Butterfly world glasshouse. On average, 69% of offspring within each family carried a B chromosome (Supplementary figure S6), suggesting that the transmission rate is not 100% which likely contributes to the B chromosome prevalence we observed.

## Discussion

Here we present the first genomic analysis of an unusual B chromosome found in a species of mealybug. This chromosome increases its transmission through males by escaping the fate that awaits other paternally-derived chromosomes under paternal genome elimination: being eliminated during meiosis. Using *de novo* generated genome data from individuals with different numbers of Bs (0-2) we were able to identify 1.37Mb of B-linked sequences (up to 7.11Mb when using more lenient criteria). These data allowed us to study both the origin and evolution of the B, as well as identify putative B-linked genes that might be involved in the drive.

### B-linked gene identity, expression, and mechanisms of drive

From the genomic analysis of the B chromosomes in *P. viburni* we confidently identified 78 B-linked genes. We then studied the expression of these genes in 2nd/3rd instar males and females as this is the development stage when meiosis takes place. We identified 13 B-linked genes that were expressed during this life stage. As the B drives during male--but not female--meiosis, we expected that any B-linked gene involved in drive to be predominantly expressed in males at this stage. We identified just one B-linked gene, g13953, that was significantly overexpressed in males compared to females and encodes a histone acetylation protein. The protein is required for the histone modification H3K56Ac which is associated with actively transcribed genes (Skalska et al. 2015) and has a role in meiosis (Wang et al. 2017). This gene is a promising candidate for its involvement in B chromosome drive in males, as the chromatin state of the B chromosome appears to play a key role in its ability to drive (Nur and Brett 1985; Nur and Brett 1988), so we expect any genes involved in chromatin remodeling could play a role. Apart from the H3K56 acetylation gene, g20108 is a gene with no known function in insects; however, it contains an integrase core domain and putative chromo domain. Even though this gene shows expression in both sexes in our RNA-seq analysis, a separate qPCR analysis shows an upregulation in male pupae compared to female last nymphal instar and earlier male stages (see Fig S8). There are also two B-linked kinesin-like protein genes, g16323 and g16324, that could be of interest. Kinesins are proteins that move along microtubule filaments and support cellular functions including meiosis. Despite the intriguing functional annotation, however, they show very low expression at this developmental stage.

In comparison with B chromosomes in other species, the B in *P. viburni* is remarkably gene rich. But, we find that most of these genes have either no or very low expression in the stage that we investigated. There are a number of possible explanations for this. First of all, it is possible that these genes are expressed in other life stages which we did not examine. Secondly, it is possible that their expression is tissue-specific and therefore challenging to detect in whole-body RNA-seq data. Finally, it is likely that the expression of B-linked genes is suppressed, especially in males when paternally-derived (which will be by far their most common state as transmission is male-biased). Throughout development in both sexes, the B appears highly condensed (heterochromatic) in comparison to the autosomes (Nur, 1966). In males, they are not only condensed but also part of the heterochromatic body at the periphery of the nucleus which contains the paternally-derived chromosomes. Recent work in another mealybug species (*P. citri*) shows that gene expression of the condensed paternal chromosomes is low but not absent (de la Filia, 2021). So even if Bs are heavily condensed, they might still show some gene expression. However, to better understand patterns of expression of B-linked genes, we will need to obtain expression data from different tissues and life stages throughout development. Combining this with population genetic analyses will also provide insights in the patterns of selection experienced by B-linked genes.

The B chromosome in *P. viburni* is not alone in its apparent ability to drive by manipulation of the chromatin state of itself and the other chromosomes. In the haplodiploid wasp *Nasonia vitripennis*, a B chromosome called *PSR* prevents the sperm chromatin from dissolving after fertilization, which leads to the elimination of all sperm-derived chromosomes with exception of *PSR* (Nur et al. 1988; Swim et al. 2012; Dalla Benetta et al. 2020). The resulting embryo develops as a haploid male rather than a diploid female, giving the *PSR* chromosome an advantage as it drives in males. The mechanism by which *PSR* accomplishes its drive through chromatin manipulation is fairly well understood: a B-linked gene called *haplodizer* appears to manipulate the histone code on the autosomes (Dalla Benetta et al. 2020). In particular, PSR induces an enrichment of H3K9Me3 and H3K20Me1, histone modifications that are associated with transcriptional suppression. Interestingly, the same histone modifications are associated with paternally-derived chromosomes in mealybugs and may play a key role in the condensation level of paternal chromosomes (Bongiorni et al. 2009; Bain 2019; Li et al. 2020). Furthermore, the degree of condensation could affect the probability that paternal chromosomes are eliminated during meiosis, with those with a more diffuse appearance more likely to pair with the maternal chromosomes and being included into functional sperm (Brown and Nur 1964; Nur and Brett 1985; Nur and Brett 1988). So a possible mechanism for drive in *P* .*viburni* would be that, similar to *N*.*vitripennis*, a B-linked gene directly manipulates the chromatin state. The two genes we have identified that have putative roles in regulating chromatin structure are therefore promising candidates for further exploration.

The mechanism by which the B chromosome drives is not likely dependent on its gene content only, but also on its structural properties, in particular its repeat content. Many B chromosomes are enriched for different types of repetitive elements. In particular satellite repeats - often unique to the B - make up a large proportion of the B chromosome in many species. For example, *PSR* in *N. vitripennis* has a repeat content of 89,8%, >70% of which consists of just 4 satellite repeats (Li et al. 2017). While the B chromosome in the migratory locust contains almost 95% repeats, with over 50% of the chromosome consisting of just a single satellite (Palacios-Gimenez et al. 2020). Although the role of B-specific repeats is still poorly understood, B-specific repeats may influence the chromatin state of the B and possibly also of the autosomes, if the B acts as a sink for chromatin proteins (Eickbush et al. 1992). Translocation experiments in *P. viburni* support this possibility, as even small fragments of the B translocated onto the autosomes decondense (and therefore presumably change their chromatin structure) during meiosis to similar degree as the entire B chromosome (Nur & Brett, 1988). This suggests it is unlikely that a single gene or region might be sufficient for controlling the Bs behaviour. In this study, we identify a number of B-specific satellite repeats that make up a significant fraction of the chromosome. These would be an interesting target for future studies into the role repeats play in the Bs ability to drive.

### Evolutionary origin of the B chromosome

Apart from determining the genetic composition of the B chromosome in *P. viburni*, we also investigated its evolutionary origin. Despite their frequent and widespread evolution, the origins of B chromosomes often remain obscure. In general, B chromosomes are assumed to have evolved from the core genome (Camacho et al. 2000). However, as they are generally rich in repetitive sequences and significantly diverged, it can be hard to pinpoint from which part of the genome they originate. There is also a suggestion that at least in some species, the B might have a hybrid origin (McAllister and Werren 1997). In *P. viburni*, it is unlikely that the B originates from pericentromeric regions of the core genome, as mealybug chromosomes are holocentric and lack a localised centromere (Bongiorni et al. 2004). We find that the B contains a large number of repetitive elements and is gene-rich. Yet, only a small number of B genes have paralogs on the core genome and several of the repeats are unique to the B. We also find that genes shared between the B and A chromosomes show substantial divergence. The same is true for homologous non-coding regions shared between the A and B chromosomes. However, when we compare divergence of B-linked sequences between the core genome of *P. viburni* and two other mealybug species, we find that they are most closely related to *P. viburni*. suggesting a within-species origin. The B chromosome does not show clear homology to any particular chromosome in the core genome and many of the genes and repeats are unique to the B chromosomes. However, many of the genes were found to be homologous to other hemipteran genes. Hence the genes unique to B chromosomes likely represent translocations from the core genome onto the B. Alternatively the B could be derived from a small portion of one of the autosomes that disassociated, something that is possible in species with holocentric chromosomes such as *P. viburni* where small chromosomal fragments can fatefully segregate.

### B chromosome frequency and transmission

We designed B-specific primers that allowed us to study the frequency of the B chromosome in two glass house populations in Edinburgh, UK. We find a high prevalence of the B chromosome in both populations; however, the B is not fixed and seems to be at about 70% frequency. B chromosome frequencies are likely determined by a number of factors. The rate at which the B drives through males is doubtless important, but so is the transmission rate through females. Previous studies suggest that the B shows a lower rate of transmission through female meiosis than expected under Mendelian inheritance (U. Nur 1962). In addition, the B may affect the fitness of its carriers; there is evidence in *P. viburni* that the B chromosome reduces male fitness and fertility (Nur, 1966). Finally, it is possible that autosomal suppressors are segregating in populations. B drive suppression mechanisms have been documented in other species, such as the post-meiotic ejection of Bs in grasshoppers(Cabrero et al. 2017)(Cabrero 2017). Nur & Brett (1987, 1988) found that the transmission rates of the B vary and are determined by the maternal genotype of a male. This suggests that there might be resistance alleles segregating in the host population. Comparing both the B chromosome chromatin landscape and autosomal gene expression in these mealybug genotypes will be another promising avenue for studying the mechanism of B drive.

## Conclusion

In this study, we identify B-linked sequences from the mealybug *Pseudococcus viburni*. This chromosome increases its transmission through males by exploiting the unusual reproductive system of mealybugs: in males all paternally-derived chromosomes are silenced (heterochromatinised) early in development and subsequently eliminated during spermatogenesis. The B chromosome is able to avoid paternal genome elimination by pairing with maternally-derived chromosomes during meiosis and ensuring its inclusion into cells destined to form mature sperm. It appears to do so by changing its chromatin state to match that of the euchromatic maternal chromosomes. We were able to identify a large number of B-linked genes that could play a role in the chromosome’s ability to drive. In particular, two genes overexpressed in males and involved in histone modification and chromatin structure are promising candidates for future work. We also identified B-specific satellite repeats that might play a role in the Bs ability to drive, either by influencing its chromatin structure, or by being the source of long or small non-coding RNA. Further studies into these B-linked genes and regions will help better understand not only the mechanisms by which the B chromosome accumulates, but also improve our understanding of the way paternally-inherited chromosomes are eliminated during male meiosis. This would provide insights into paternal genome elimination, a peculiar but widespread reproductive system that has evolved repeatedly across arthropods.

## Materials and methods

### Mealybug populations and cultures

The obscure (or glasshouse) mealybug *Pseudococcus viburni* (Hemiptera: Pseudococcidae) is a small phloem-feeding insect with a global distribution. It is able to feed on a large number of plant species and is a common pest across many regions in glasshouses, fruit orchards and vineyards. We collected egg-laying *P. viburni* females from a variety of host plants in the glasshouses of the Royal Botanic Gardens of Edinburgh (UK) and transferred them to sprouted potatoes. These experimental populations were kept at 25°C degrees (16h-light/8h-dark photoperiod without humidity control) in tupperware boxes modified to have a large hole on the lid and sealed with a screen printing mesh, with a tissue paper at the bottom of the box to absorb excess moisture.

Species identification was confirmed using multiplex PCR on extracted DNA from single individuals per collected population, using primers previously published (Daane et al. 2011). gDNA was extract using the prepGEM Insect kit (ZyGEM, New Zealand) following manufacturer instructions, and multiplex PCR was carried out using the Qiagen Multiplex PCR Kit (Qiagen), following reaction specifications in (Daane et al. 2011). 4ul of PCR product was visualised by electrophoresis on a 2% agarose gel stained with GelRed Nucleic Acid Gel Stain (Biotium).

We established the experimental lines used in this study in plastic fly medium bottles containing a sprouted potato by transferring single egg-laying females and letting the offspring grow, reach adult stage and mate. Several lines were established from the base population and these were regularly screened for B chromosomes using karyotyping. Information on the final lines used for this study can be found in Supplementary Table S1. Successive generations were established by repeating the process once egg-laying female offspring could be spotted. This resulted in a number of highly inbred iso-female lines.

### B chromosome screening

To screen for B chromosomes, we prepared cytological slides at embryonic stage (1-2 days after egg laying) and during male second instar and prepupal stages. For embryos, laid eggs were transferred onto a coverslip with 20uL of 45% acetic acid, and squashed using a microscope slide. The coverslip was then snapped after emerging the microscope slide in liquid nitrogen, and about 40 uL of 1:3 acetic acid:ethanol was applied on the tissue and air dried. For male juvenile stages, whole males were fixed in Bradley Carnoy between 24-72 hours at 4°C. Testes were then dissected on a coverslip in 20uL of 45% acetic acid. A slide was then applied on top of the coverslip and turned coverslip face up for squashing between a piece of filter paper. The coverslip was then snapped by dipping the slide into liquid nitrogen. 40uL of Ethanol:Acid Acetic 3:1 was applied on the sample and the slide was left to dry. DNA was visualised with Vectashield Antifade Mounting Medium with Dapi (Vector Laboratories), and images were taken on a Leica DM2500 fluorescence microscope, using the Leica application suite software. Link to detailed protocol: dx.doi.org/10.17504/protocols.io.mnec5be.

### Sequencing of B and non-B experimental lines

#### DNA-seq

We transferred 1-3 egg-laying females from two B-carrying experimental lines (PV04 and PV13) and two non-B lines (PV21 and PV23) from the cultures to individual bottles and let them lay eggs for 24h. The females were then flash frozen and stored at -80C. After karyotyping of eggs to confirm expected presence/absence of B chromosomes, we extracted genomic DNA from a pool of three females from the PV13 line and single females from the other three lines using a combination of two DNA extraction kits. The full protocol is available online (dx.doi.org/10.17504/protocols.io.pvgdn3w). The amount of extracted DNA was measured using the Qubit dsDNA Assay Kit (ThermoFisher). PV04, PV13, PV21 and PV23 TruSeq DNA Nano libraries (Illumina) with 350bp insert sizes and an additional 550bp library for PV13 were generated by Edinburgh Genomics (UK) and sequenced on a Novaseq S1 flowcell to yield 203-402M 150bp paired-end reads per line. Additionally, a PacBio SMRTbell 20Kb library (Pacific Biosciences) was generated from a pool of 60 adult PV13 females and sequenced on a SMRT Cell 1M v3 on the PacBio Sequel platform, generating 28.6Gb of data (number of subreads = 1.7M, subread N50 = 14.9Kb). All reads were subjected to filtering for endosymbionts that are not part of this study and deposited to ENA under accession (PRJEB47083).

#### RNA-seq

We isolated, sexed, and flash-froze pools of 30-60 individuals from PV04, PV13, PV21, and an additional experimental line, PV15, from the same original non-B population. All specimens were late 2nd or early 3rd instars. This is the stage when males spin a cocoon and undergo a process similar to metamorphosis and therefore the earliest stage at which specimens can be sexed reliably. We extracted total RNA from 3-4 biological replicates per sex and line with a PureLink RNA purification kit (Thermo Fisher Scientific, USA) and validated the samples with the Bioanalyzer RNA 6000 Nano kit (Agilent). 23 TruSeq stranded mRNA-seq libraries (Illumina) were generated by Edinburgh Genomics and sequenced on the NovaSeq platform, yielding 86-136M 50bp paired-end reads per sample.

### Genome assembly and annotation

We assembled the PacBio subreads with wtdbg2/Redbean v2.5 (Ruan and Li 2020) using the default settings for PacBio Sequel technology. To polish the raw assembly, we ran three rounds of HyPo v1.0.3 (Kundu et al. 2019) using the PacBio reads and Illumina short reads from the 350 and 550bp PV13 libraries (see below for read processing). During polishing, we used minimap2 v2.17 (Li 2018) to map long and short reads to the successive assemblies with the --secondary=no setting.

After polishing, we performed a first round of scaffolding using transcriptomic evidence with SCUBAT2 (https://github.com/GDKO/SCUBAT2), using a *de novo* transcriptome assembly assembled from all male and female RNA-seq samples using Trinity-v2.8.5 (Grabherr et al. 2011). We only used transcripts with TPM>2 in all 23 RNA-seq samples and a maximum intron size of 50,000. We then ran a second round of scaffolding with BESST v2.2.8 (Sahlin et al. 2014) and the Illumina short reads. Then, we followed the blobtools v1.1.1 pipeline (Laetsch and Blaxter 2017b) to remove contaminants from the assembly: briefly, contigs/scaffolds with homology hits to Proteobacteria and increased genomic coverage, indicating a bacterial endosymbiont origin, and contigs without taxonomic hits and coverage <2). Throughout the process, we used BUSCO v4.0.6 (Simão et al. 2015) with the Hemiptera and Insecta gene sets to assess completeness of the successive iterations of the assembly.

To annotate the genome, we first ran RepeatMasker v4.1.0 (Smit et al. 2019) to soft mask repetitive regions in the assembly using a repeat library combining a de novo library built with RepeatModeler v2.0 and the hemipteran RepBase database. To predict genes, we followed the BRAKER2 v2.1.4 (Hoff et al., 2015) pipeline using our 23 RNA-seq samples, which we mapped to the assembly using STAR v2.7.4a (Dobin et al. 2013) in the single pass mode. We annotated the predicted transcripts with homology-based searches against the UniProtKB/Swiss-Prot (BLASTp v2.9.0, E-value = 1e-25) and reference proteome databases (diamond v0.9.29, E-value = 1e-25) and running InterProScan (Quevillon et al. 2005). to assign Pfam and InterPro domains and GO terms.

### Identification of B chromosome scaffolds in the assembly

We took three complementary approaches to define a set of B-linked scaffolds in the assembly: genomic coverage, assembly mapping, and kmer coverage.

To identify candidate B regions based on differential genomic coverage, we mapped the Illumina short reads generated for the four experimental lines to our reference genome, which should contain the B. We used FastQC v0.11.5 (https://qubeshub.org/resources/fastq) for read quality control and fastp v0.20 (Chen et al. 2018) for quality and adapter trimming with the following settings: --detect_adapter_for_pe --cut_by_quality5 --cut_by_quality3 --cut_window_size 4 --cut_mean_quality 20 --trim_poly_g. We mapped the reads from each line to the reference genome using bwa mem v0.7.17-r1188 (Li 2013) with default settings (mapping rates 94.8-97.8%) For PV13, we first mapped both 350bp and 550bp libraries separately and then merged the resulting BAM files. We then used samtools v1.9 (Li et al. 2009) to filter out secondary mappings and bamtools (Barnett et al. 2011) to remove reads with edit distance >0 (NM tag) in order to keep only primary mappings with no mismatches. We collected numbers of reads mapping to each scaffold using samtools idxstats and estimated scaffold coverage differences between pairs of lines after normalising by the median coverage difference across all scaffolds. We defined two thresholds to identify putative B-linked scaffolds: a ‘relaxed’ threshold of log2(B+ line/B-line)>0.58 in all 4 pairs (which considers a hypothetical B region of autosomal origin that is present three times in an individual carrying one B chromosome and twice in a non-B individual) and a more conservative threshold of log2>2 (the ‘strict’ criterion).

To identify B scaffolds via assembly mapping, we assembled the trimmed short read libraries separately into assemblies for each of the four lines with SPAdes v3.11.1 (Bankevich et al. 2012) with the following settings: --only-assembler -k 21,33,55,77,99,127. Then, we mapped the four assemblies to the reference genome using the nucmer aligner from MUMmer v4.0.0beta2 (Marçais et al. 2018) and used the dnadiff wrapper to extract 1-to-1 alignments between the reference genome and the assemblies and determine which reference scaffolds map only to B+ lines.

To identify B scaffolds using kmer coverage, we loosely followed the strategy developed by Hodson et al. (2021). This method uses differences in k-mer frequencies between sequencing libraries to identify overrepresented (and thus putatively B-linked) kmers in B+ libraries independently of genome assembly. We used KMC3 (Kokot et al. 2017) to extract 27-mers with at least 3 counts in all libraries and their coverages from the raw reads (after removing contaminant reads using the blobtools pipeline; see above). We used total kmer coverage estimates from each sequencing library (as determined with GenomeScope v1, http://qb.cshl.edu/genomescope/info.php) to account for differences in sequencing depth between lines and normalise coverages of individual kmers. Since B chromosome kmers may be unique to Bs or shared between A and B, we used the relaxed threshold of log2(B+/B-lines)>0.58 (see above) to assign putative B origin to kmers. Considering only kmers with normalised coverage >5 in at least one line, we classified 49.7M kmers as putatively B-linked, while 504M kmers were considered autosomal. We then mapped these kmers to the assembled scaffolds in the reference genome using bwa mem with the following settings: -k 27 -T 27. Scaffolds with as many putative B as autosomal kmer mappings (log2>=0) were considered B candidates.

### Gene expression quantification and differential expression analyses

We used FastQC and fastp for raw RNA-seq data processing and quantified gene expression levels in all 23 samples using RSEM v1.3.0 (Li and Dewey 2011) with the STAR aligner (Dobin et al. 2013).

We evaluated differential gene expression using the edgeR/limma workflow described in (Law et al. 2016). For the sex-biased differential expression analysis, we only included sex as a factor. Briefly, we removed lowly expressed genes (minimum read count = 10), converted raw data to log2(CPMs) and normalised expression distributions with the TMM method using edgeR (Robinson et al., 2010). We then validated all our RNA-seq samples using density plots, log2(CPM) boxplots, MDS plots and correlation heatmaps. We transformed the data using the voom function and built a linear model in limma with empirical Bayes moderation (Ritchie et al., 2015). We required a threshold of log(fold change)>1 (twice expressed in one sex compared to the other) for differentially expressed genes (with an FDR<0.05). We conducted the differential expression analysis between B+ and B-samples identically, with sex and B presence as factors, but with the minimum read count and log(fold change) lowered, respectively, to 5 and 0.58 (50% higher expression) to account for the overall low expression of B1 genes (Supplementary Figure S3). We conducted gene ontology (GO) enrichment analyses for differentially expressed using the hypergeometric test implemented in the GOStats R package (Falcon and Gentleman 2007) with a p-value cutoff of 0.01.

### PCR amplification of B-linked primers

We used Primer3 (https://primer3.ut.ee) to design PCR primers for the longest B-linked scaffold, scaffold_552 (Supplementary Table S3). For a single individual, PCR amplification was performed in a 25 μl reaction volume containing 1 μL of prepGEM-extracted gDNA product, 12.5 µl DreamTaq Green PCR Master Mix (ThermoFisher), 0.25 µl forward primer and 0.25 µl reverse primer. PCR reactions were performed under the following conditions: initial denaturation at 95°C for 1 min, 30 cycles of denaturation at 95°C for 30s, annealing at 55°C for 30s and extension at 72°C for 1m and a final extension step at 72°C for 5 minutes. PCR products were placed under electrophoresis at 100V in a 2% agarose gel with 1% TAE buffer for 40 minutes with a 1kb DNA ladder.

## Acknowledgements

We would like to thank Patrick Ferree for his comments, help and support designing this study, glasshouse staff from the Royal Botanical Gardens and Butterfly House Edinburgh for allowing us to collect specimens. The work was funded by an Marie Skłodowska-Curie Individual Fellowship to IV, a Dorothy Hodkins Fellowship to LR while KJ and AM were funded by an ERC starting grant.

